# Engineering One-Signal Activated TILs with CD3ζ-CD28 Chimeras Optimizes the Antitumor Efficacy of TIL-Based Immunotherapy

**DOI:** 10.1101/2024.03.21.586203

**Authors:** Yongqiang Wu, Xueshuai Ye, Yufeng Zhu, Shengchao Liu, Jiabao Song, Yutong Zhang, Qichen Yuan, Fei Liu, Dejing Kong, Lianmei Zhao

**Affiliations:** Gene Editing Research Center, Hebei University of Science and Technology, Shijiazhuang, 050000, Hebei, China; Research Center, The Fourth Affiliated Hospital of Hebei Medical University, Shijiazhuang, 050017, Hebei, China; The Affiliated Hospital and School of Clinical Medicine, Hebei University of Engineering, Handan, 056000, Hebei, China; College of Food Science and Biology, Hebei University of Science and Technology, Shijiazhuang, 050000, Hebei, China; Department of Chemical and Biomolecular Engineering, Rice University, Houston, 77005, Texas, USA

## Abstract

Adoptive T cell therapy (ACT) is a promising therapeutic option that enhances the anti-cancer immune response to recognize and eliminate tumor burden. Recently, ACT modalities with autologous tumor-infiltrating lymphocytes (TILs) have shown robust anti-tumor efficacy in solid tumors. However, this therapeutic strategy’s application remains suboptimal in stability and faces critical challenges, primarily due to the lack of costimulatory signals. Here, we unveil a novel engineering strategy one-signal-activated TIL (OSAT), which integrates CD3ζ with secondary signaling domains, to bypass the classical dual-signal requirement for T cell activation. We demonstrate that OSAT molecules 2 (CD3ζ-CD28, M2) and 7 (CD3ζ-CD137-CD28, M7) mediate efficient mono-signal activation, and recapitulate both the activation signatures and functional markers of mock T cells stimulated with dual signals. Notably, OSAT exhibited enhanced cytotoxicity and *in vivo* anti-tumor activity across syngeneic tumor models, especially when combined with PD-1 blockade therapy. Mechanistically, OSAT can be fully activated by first signal stimulation, leading to upregulation of activation and metabolic reprogramming genes, production of IL-2/IFN-γ/TNF-α, initiation of cytotoxicity, and enhanced TIL antitumor efficacy. These findings unveil integrating CD3ζ with secondary signaling domains as a higher efficacy modification for TILs with therapeutic potential for clinical cancer immunotherapy.

## Introduction

Cancer immunotherapies including ACT and immune checkpoint inhibitor (ICI) have revolutionized cancer treatment^1–4^. TIL therapy is one of the most important ACT strategies, which involves the isolation and purification of specific antitumor lymphocytes from solid tumor tissue, followed by their amplification and activation before administration^5,6^. A key advantage of TILs lies in their inherent capacity to target patient-specific tumor antigens and infiltrate tumor tissues, thereby enabling the elimination of tumor cells in the therapeutic setting of solid malignancies^7,8^. Nevertheless, TIL therapy still faces obstacles in the treatment of solid tumors, as the diverse microenvironments and immune cell states result in its inability to efficiently kill tumor cells^9^. Specially, the inefficient tumor-killing capacity of TILs in solid tumors is closely linked to defects in T cell activation signaling^5,6,10^.

The efficient anti-tumor function of TILs relies on intact T cell activation signals. T cell activation follows the classical “two-signal model” that is TCR-CD3 engagement with MHC-bound peptides and CD28 co-stimulation by antigen-presenting cells (APCs) ^10,11^. The first signal is MHC class I molecule-antigen peptide complexes that is abundant in tumor tissues, and could be specifically recognized by TCRs. For the second signal CD28 co-stimulation, however, the function of which is often impaired by immunosuppressive factors and regulatory immune cells in the tumor microenvironment (TME), resulting in insufficient co-stimulation and ultimately T-cell anergy^12^. Consequently, T cells may remain inactivated due to the absence of co-stimulatory signals from APCs, leading to immune tolerance^13^. Moreover, sustained CD28 co-stimulation is essential for the self-renewal and differentiation of TCF-1⁺ PD-1⁺ CD8 T cells, as well as for maintaining the functionality of effector T cells^13,14^. Thus, these aberrant signaling states culminate in T cell dysfunction, compromising anti-tumor immunity^15,16^. However, combining TIL with CD28-activating antibodies has not been a viable strategy to date, thus alternative strategies to deliver CD28 co-stimulation in a tumor antigen-specific manner are urgently needed. For instance, in the 2006 TGN1412 phase I clinical trial involving CD28 monoclonal antibodies, six healthy male volunteers rapidly developed severe systemic inflammatory responses after receiving a single dose of the antibody, resulting in critical conditions^17^. This demonstrates that therapeutic use of CD28 monoclonal antibodies can elicit significant adverse effects, posing substantial challenges for their application in augmenting TIL-mediated tumor combat. Given the risks of systemic toxicity with CD28 antibodies, alternative strategies to deliver CD28 co-stimulation in a tumor antigen-specific manner are needed.

The TCR-CD3 complex is a multiprotein assembly consisting of the TCR-αβ heterodimer, together with the signaling components composed of the ζζ homodimer and CD3δ, CD3ε, and CD3γ subunits^18^. Each CD3δ/ε/γ subunit contains an extracellular immunoglobulin domain, a transmembrane region, and a cytoplasmic intracellular domain that harbors a single immunoreceptor tyrosine-based activation motif (ITAM)^19^. Notably, the CD3ζ subunit differs structurally and functionally. It has a short extracellular stalk and a long ICD containing three ITAMs, enables it to amplify signaling more efficiently than other CD3 subunits^18,19^. Furthermore, this superior signaling amplification is pivotal for driving potent and sustained T-cell responses against pathogens or aberrant cells. The clinical success of CAR-T therapy provides direct evidence that fused intracellular activation domains (ICD) of CD3ζ and CD28/137 are sufficient to fully activate T cells^5–6^. Nevertheless, translating such success to the clinical context of solid tumors remains a major therapeutic challenge^20^. Recent advances in CAR-T engineering for solid tumors have focused on addressing antigen heterogeneity via bispecific/tandem CARs targeting multiple antigens, enhancing persistence through CD137 costimulation, or combining with checkpoint inhibition to achieve synergistic effects^21,22^. However, antigen heterogeneity is a common driver of tumor escape, while desmoplastic stroma and immunosuppressive molecules in the TME greatly limits the infiltration of infused CAR-T cells into tumor sites^23,24^. These barriers underscore the need for alternative engineered T-cell therapies that balance antigen specificity, heterogeneity, TME adaptability, and safety. Thus, there is a paucity of research on the expression of CD3ζ fused with costimulatory factors in TILs, which represents a critical unaddressed area for the development of next-generation adoptive T-cell therapies for solid malignancies.

Herein, we introduce the OSAT approach, intracellularly linked fusion proteins of full-length CD3ζ and CD28/137 intracellular activation domains with endogenous TCR, to optimize the T cell activation process to enhance the antitumor efficacy of TILs. Our innovation involves the development of a single-step activation receptor that integrates essential intracellular co-stimulatory signals within TIL cells. Moreover, we propose that the modified TIL cells can directly identify and eliminate tumor cells through bypassing the need for co-stimulatory signals. Importantly, these modified TILs retained their specific tumor recognition and tissue infiltration capabilities, and no significant inflammatory responses were observed *in vivo*. Notably, we validated the efficacy and safety of the modified TILs in combination with PD-1 blockade in syngeneic tumor models, providing a strategy for enhancing the efficacy of TILs in combination with PD-1 blockade. Taken together, this study innovatively develops the OSAT approach to overcome traditional TILs’ reliance on external co-stimulatory signals, and provides an effective, safe therapeutic strategy to advance TIL-based immunotherapies and address clinical challenges in treating malignant solid tumors.

## Results

### OSAT design and screening of OSAT molecules

The clinical success of CAR-T therapy directly validates that the structurally and functionally distinct CD3ζ subunit can enable full T cell activation when fused with CD28/137-ICD^5,6^. Thus we hypothesize that ζ subunit could rejuvenate exhausted TILs and augment anti-tumor responses through streamlined yet robust signaling. To address this, we engineered OSAT molecules consisting of CD3ζ fused with ICD of costimulatory receptor (Fig. 1a). Then we designed OSAT molecules by fusing full-length CD3ζ with ICD of CD28 and/or CD137 in combinatorial configurations (Fig. 1b; Supplementary File). To enable screening, *EF1α* promoter-driven lentiviral vectors expressing CD3ζ-CD28/CD137 chimeras with enhanced green fluorescent protein (EGFP) were constructed and transduced into Jurkat cells. The stable cell lines were established via puromycin selection and the expression of OSAT molecules were successfully detected by flow cytometric analysis (Fig. 1c). The functional validation of OSAT constructs was initially determined by the secretion of cytokines (IL-2/IFN-γ/TNF-α). Accordingly, OSAT molecule 2 (CD3ζ-CD28, M2) and 7 (CD3ζ-CD137-CD28, M7) exhibited significant cytokine secretion (Fig. 1d). Next, we characterized the expression changes of activation markers (CD25 and CD69) in Jurkat cells upon 24 hours stimulation with anti-CD3 or control IgG. Compared to mock Jurkat cells, OSAT-engineered Jurkat cells exhibited a significant upregulation of these activation markers upon anti-CD3 stimulation (Fig. 1e). Taken together, these results demonstrated that novel engineering constructs M2 and M7 possess the potential to activate T cells via mono-signal pathway, and they were therefore selected for subsequent downstream validation.

**Fig. 1.**
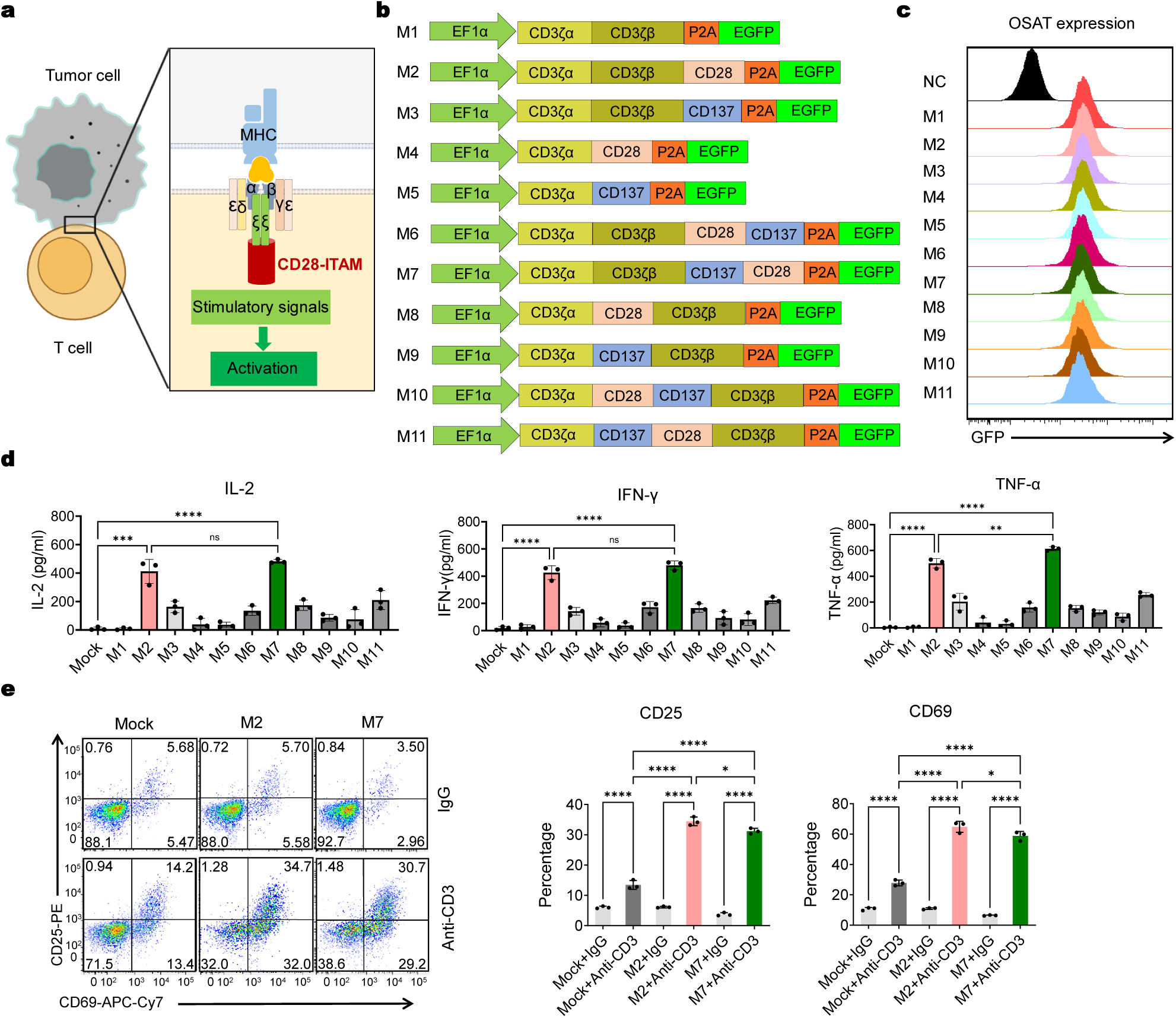
Schematic representation of T cell activation mechanisms and screening of OSAT molecules. **(a)** Hypothetical working model of OSAT: The chimeric construct integrates CD3ζ with CD28 costimulatory domains, enabling antigen-independent T cell activation upon tumor cell contact. **(b)** Molecular architecture of OSAT constructs (CD3ζα: N-terminal extracellular/ transmembrane domain (amino acids 1-51), CD3ζβ: C-terminal ITAM-containing cytoplasmic domain (amino acids 52-163), CD28: CD28 intracellular ITAM domain, CD137: CD137 intracellular ITAM domains). **(c)** Flow cytometric analysis the OSAT expression. **(d)** Cytokines secreted were measured 24 h post-stimulation. **(e)** Expression of activation markers of Jurkat cells after 24 h of stimulation with anti-CD3 (1 μg/mL) or IgG (1 μg/mL). Data represent mean ± SEM (n = 3 biologically independent experiments). Statistical significance was determined by one-way ANOVA with tukey’s multiple comparisons test (*\*P* < 0.05, *\*\*P* < 0.01, *\*\*\*P* < 0.001, *\*\*\*\*P* <0.0001; ns = not significant).

### T cell activation mediated by OSAT molecules M2 and M7 under mono-signal stimulation

T cell activation via TCR engagement and costimulatory receptor engagement initiates membrane depolarization, triggering Ca²⁺ release from the endoplasmic reticulum into the cytosol^19^. Elevated cytosolic Ca²⁺ levels activate downstream signaling cascades, including the nuclear factor of activated T cells (NFAT) pathway, which coordinately enhances transcription factor activity (NF-κB, AP-1) critical for T cell proliferation, differentiation, and effector function programming^25–27^. Next, to characterize OSAT mediated signaling, we leveraged Jurkat cells, a tractable model system with reduced biological variability compared to primary cells. Our result showed that mono-signal anti-CD3 stimulation of OSAT-engineered Jurkat cells (OSA-Jurkat) induced sustained Ca²⁺ flux comparable to dual-stimulated mock Jurkat cells (Fig. 2a). Moreover, phospho-protein analysis revealed significant upregulation of activated AP-1 and NF-κB in M2/M7 OSA-Jurkat cells 1 hour post-stimulation, whereas mock Jurkat cells showed no detectable activation under mono-signal conditions (Fig. 2b).

**Fig. 2.**
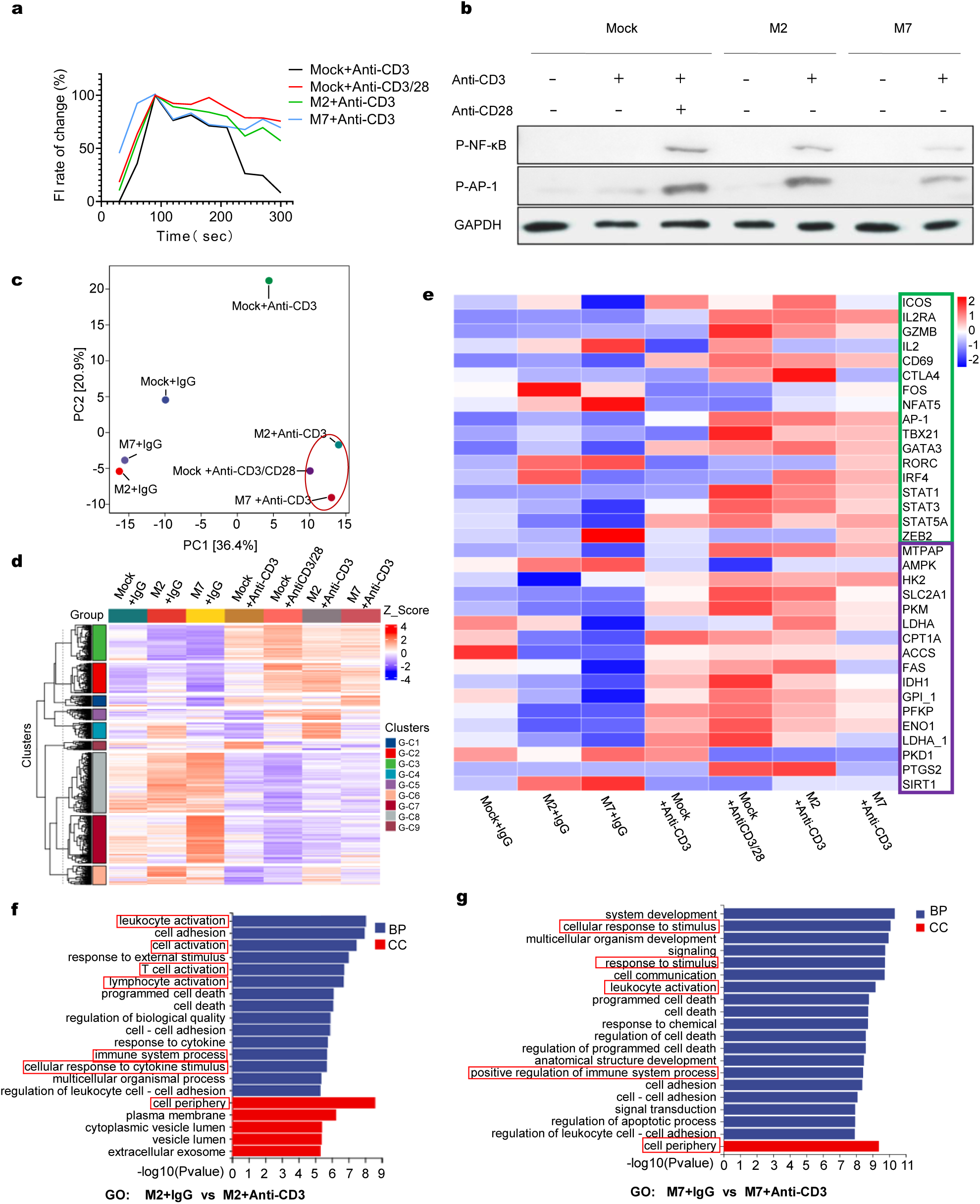
OSAT molecules recapitulate dual-signal activation programs under mono-signal stimulation. **(a)** Intracellular Ca²⁺ dynamics in Jurkat cells measured by Fura-8 AM-based calcium imaging following anti-CD3 (1 μg/mL) and/or anti-CD28 (1 μg/mL) stimulation. Data represent mean ± SEM (n = 3 biologically independent experiments). **(b)** Immunoblot analysis of phosphorylated transcription factors in Jurkat cells post-activation. Cells were stimulated with anti-CD3/CD28 for 60 minutes and probed with antibodies against phospho-NF-κB p65 (Ser536) and phospho-AP-1 c-Jun (Ser63). GAPDH served as a loading control. **(c)** PCA plot from the full transcriptome of mock-Jurkat or OSA-Jurkat cells following indicated stimulations. **(d)** Heatmap of differentially expressed gene clusters of CD3 activations. Clusters highlight distinct transcriptional responses to mono-vs. dual-signal stimulation (Clustering was determined using k-means). **(e)** Heatmap of Z-score normalized expression for T cell activation markers (green) and metabolic pathway genes (purple) in OSA-Jurkat and mock Jurkat cells 12 hours post-stimulation. **(f-g)** GO enrichment analysis comparing M2 + IgG vs. M2 + Anti-CD3 and M7 + IgG vs. M7 + Anti-CD3 Jurkat cells. The top 20 most significantly enriched GO terms for differentially expressed genes are shown. GO enrichment analysis was performed using the cluster Profiler R package. Differentially expressed genes were annotated to GO terms, and enrichment was calculated *via* hyper geometric distribution.

Next, we performed transcriptomic profiling to verify the functional equivalence between OSA-Jurkat and dual-stimulated mock Jurkat cells. Principal Components Analysis (PCA) of the transcriptome data showed that OSA-Jurkat cells with mono-stimulation clustered with dual-stimulated mock-Jurkat cells (Fig. 2c). Global gene expression patterns also exhibited high similarity between OSA-stimulated Jurkat cells (mono-stimulation) and mock-stimulated Jurkat cells subjected to dual stimulation (Supplementary Fig. 1a). Moreover, venn analysis of differentially expressed genes (DEGs) identified about 30% overlap between OSA-Jurkat and dual-signal activation, with divergent enrichment in co-stimulatory and epigenetic regulatory pathways (Supplementary Fig. 1b). Furthermore, hierarchical clustering revealed the activation of OSA-Jurkat cells under mono-signal conditions clustered closer to dual-stimulated mock Jurkat cells, suggesting that the mono-signal stimulation of OSA-Jurkat cells is sufficient to recapitulate the functional equivalence of dual-signal activation in mock Jurkat cells, consistent with their convergent transcriptional profiles and clustering patterns (Fig. 2d). Notably, functional annotation of DEGs highlighted upregulation of activation genes (such as ICOS, IL2RA, GZMB) and metabolic reprogramming genes (such as MTPAP, HK2, PKM) in both dual-stimulated mock cells and mono-stimulated M2-OSA cells, highlight the important role of OSA in driving T cell functional activation (Fig. 2e). We next performed Gene Ontology (GO) enrichment analysis to further validate signaling equivalence, and the results demonstrated that both dual-stimulated mock Jurkat cells and OSA-Jurkat cells exhibited statistically significant enrichment in T cell activation-associated biological processes (BP) and cellular components (CC) (Fig. 2f-g; Supplementary Fig. 1c). In contrast, mono-stimulated mock Jurkat cells showed no such enrichment (Supplementary Fig. 1d). These results illustrate that dual-signal stimulation and OSA-mediated mono-signal activation in Jurkat cells is sufficient to elicit the transcriptional enrichment of biological processes and cellular components associated with T cell activation, whereas mock Jurkat cells fails to induce this specific activation-related GO signature. Collectively, we demonstrate that OSAT molecules recapitulate dual-signal activation programs under mono-signal conditions through combinatorial co-stimulatory signaling.

### Functional validation of OSA-Engineered primary T cells with optimized gene delivery

To enable OSA-mediated modification of TILs for tumor eradication, a safe, efficient, and rapid gene delivery method is critical. For instance, the Cas9-HITI gene editing method precisely inserts the OSA sequence into the CD3ζ locus while knocking out wild-type CD3ζ, minimizing competition between endogenous CD3ζ and OSAT molecules^28^. Thus, we initially evaluated multiple single-guide RNAs (sgRNAs) targeting the CD3ζ locus and identified the most effective sgRNA *via* T7E1 analysis (Supplementary Fig. 2a-c). We designed an OSA-donor vector (incorporated a 2A-GFP cassette into the C-terminus) and co-transfected primary T cells with Cas9-sgRNA and OSA-Donor *via* electroporation using this sgRNA. At 72 hours post-transfection, the optimal sgRNA achieved about 14% editing efficiency, whereas lentiviral transduction of primary T cells yielded about 64% efficiency (Fig. 3a). To explore transient high-level expression, we employed a self-replicating plasmid with S/MAR ori^29^. Moreover, long-term stability analysis revealed that gene editing and lentiviral methods maintained stable OSA expression over two weeks, whereas electroplasmid transfection showed reduced expression (Supplementary Fig. 2d-f). Subsequently, comparable effects on T cell proliferation indicate that lentiviral transduction offers superior stable expression efficiency (Fig. 3b).

**Fig. 3.**
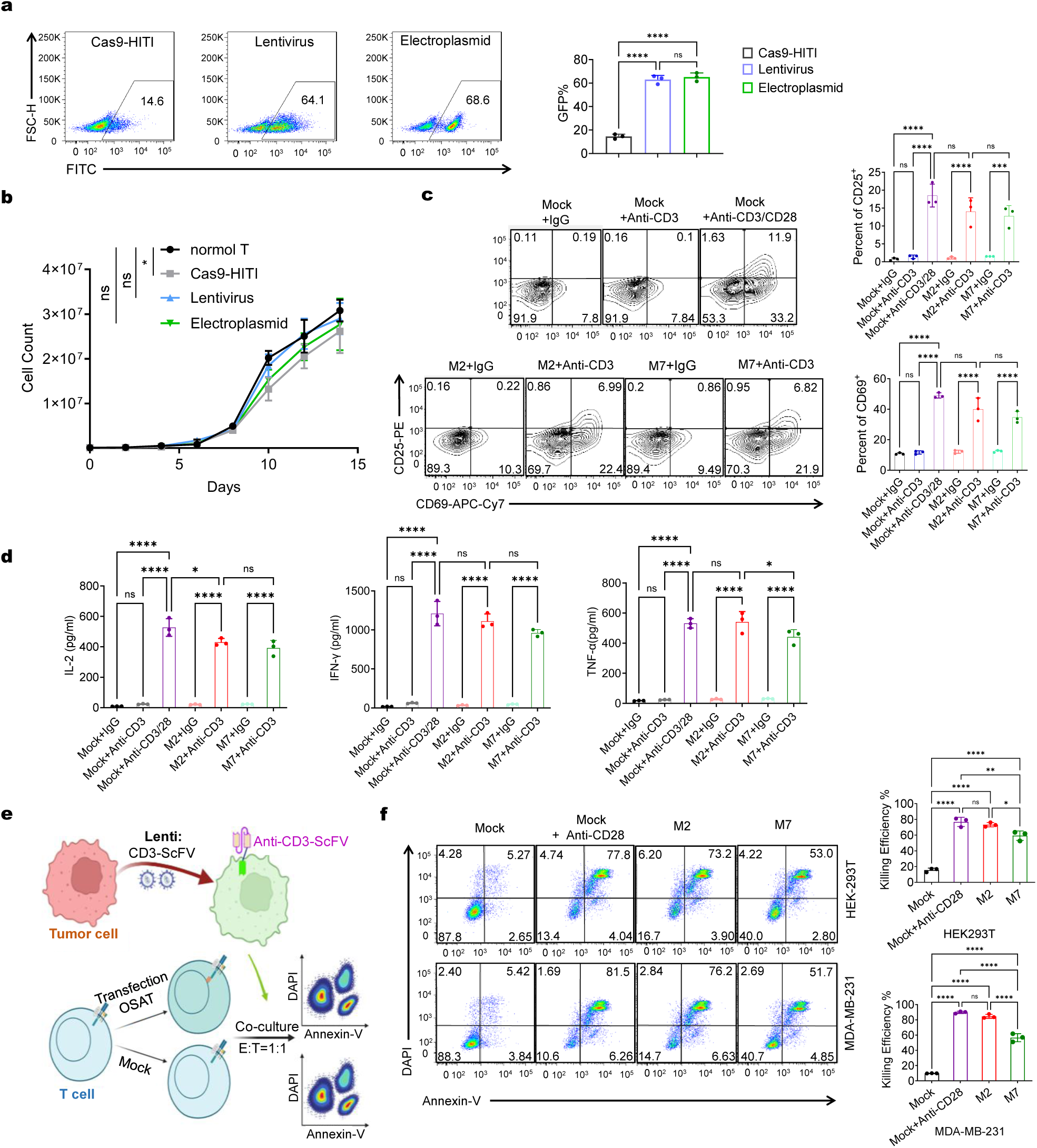
Functional validation of OSAT molecule efficacy in primary T cells. **(a)** Flow cytometry analysis of OSA expression efficiency 72 hours post-transduction. Left: Representative flow plots showing OSA expression in primary T cells under indicated conditions. Right: Quantification of OSA-positive cell populations across three independent experiments. **(b)** Proliferation curves of unmodified T cells, lentivirus-transduced OSA-T cells, Cas9/HITI-engineered OSA-T cells, and electroporated control T cells. Data pooled from three donors. **(c)** T cell activation profiling. Left: Flow cytometry gating for CD25 and CD69 expression under specified stimulation conditions. Right: Quantification of CD25^+^/CD69^+^ populations across triplicate experiments. **(d)** Cytokine secretion profiles measured by ELISA. IL-2, IFN-γ, and TNF-α concentrations in supernatants following gene transfer and stimulation under indicated conditions. **(e)** Experimental workflow for assessing OSA-T cell cytotoxic activity mediated solely by primary (CD3) signaling. **(f)** Cytotoxic efficacy of primary T cells against HEK293T and MDA-MB-231 cell lines. Killing efficiency (%) was evaluated using a dual-marker (Annexin-V and DAPI) flow cytometry assay combined with quantitative analysis. All graphs represent triplicate experiments (n = 3 donors). Error bars denote mean ± SEM. Statistical significance was assessed by one-way ANOVA with tukey’s multiple comparisons test for b-d and f (*\*P < 0.05, **P < 0.01, ***P < 0.001, ****P <0.0001*; ns = not significant).

To evaluate OSAT functionality in primary T cells, we transduced M2/M7 constructs into human primary T cells using lentiviral vectors and performed functional assays under indicated stimulation. Flow cytometry showed minimal CD25/CD69 in unstimulated or anti-CD3 activated mock T cells, whereas OSA-engineered T cells exhibited robust upregulation of these markers upon anti-CD3 stimulation alone (Fig. 3c). Moreover, cytokine secretion profiles (IL-2, IFN-γ, TNF-α) demonstrated significant increases in all activated groups relative to controls, with no statistically significant differences between M2-OSA-mediated and dual-signal activation (Fig. 3d). Furthermore, functional validation was utilized to evaluate the cytotoxic ability with HEK-293T/MDA-MB-231 cells engineered to express CD3-specific ScFV on their surface (Fig. 3e). Accordingly, Mock T cells required CD28 co-stimulation for consistent target cell killing, whereas OSA-modified T cells exhibited potent cytotoxicity without CD28 engagement (Fig. 3f). Taken together, these results confirm that OSAT molecules fully activate primary T cells through TCR/CD3 engagement alone, bypassing traditional co-stimulation requirements while retaining cytotoxic effector functions.

### OSAT enhances antitumor functionality in multiple tumor models *in vitro*

To further validate the antitumor efficacy of OSAT *in vivo*, human OSAT molecules sequences were substituted with murine orthologs and evaluated in the EL-4 murine T cell line. Following electroporation with OSAT molecules or negative control (Mock), cells were stimulated with anti-mouse CD3 monoclonal antibody 72 hours later. Accordingly, OSA-modified EL-4 cells demonstrated significantly elevated IL-2 and IFN-γ secretion (Fig. 4a, b), confirming functional conservation of OSAT molecules across species. Subsequently, subcutaneous tumor models using B16, MC38, and TC-1 cell lines were generated to assess OSAT functionality. Upon reaching about 0.5 cm³ tumor volume, CD3⁺ TILs were isolated from dissociated tumors *via* magnetic-activated cell sorting and transfected with OSA constructs (Fig. 4c). Accordingly, co-culture assays with autologous tumor cells at varying effector-to-target (E:T) ratios revealed significantly enhanced cytotoxicity by OSAT compared to non-modified TILs across all tested ratios (Fig. 4d). In contrast, PBMC-derived CD3⁺ T cells showed no tumor-specific killing activity, confirming TILs’ intrinsic targeting specificity (Fig. 4d-e). ELISA analysis further demonstrated significantly higher secretion of IL-2 and IFN-γ from OSAT at a 20:1 E:T ratio compared to controls (Fig. 4e). Collectively, these results indicate that OSAT exhibit enhanced cytokine secretion capacity and high killing efficiency *in vitro*.

**Fig. 4.**
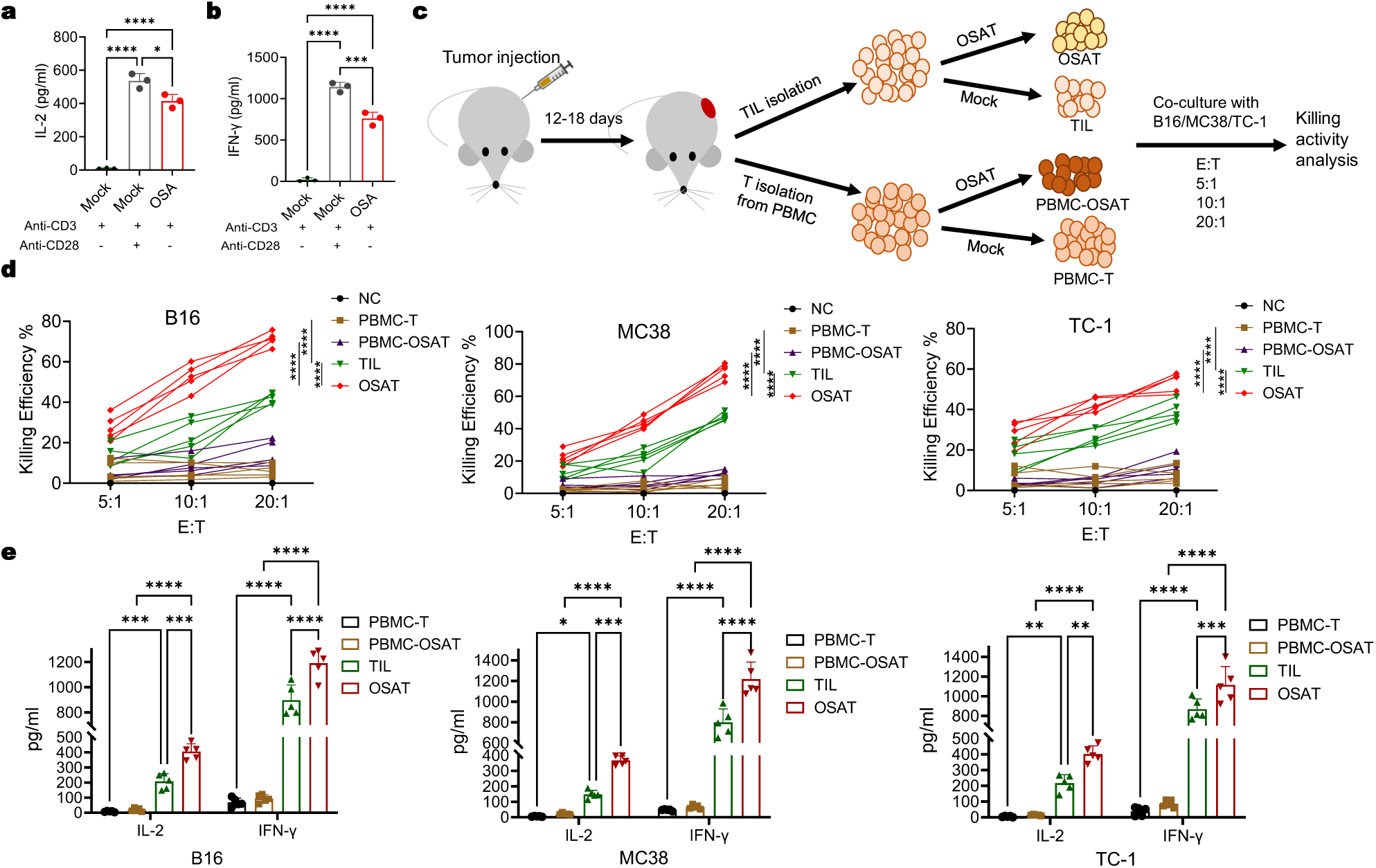
OSAT is efficacious against multiple tumors *in vitro*. **(a-b)** Cytokine production analysis of IL-2 and IFN-γ in EL-4 mouse T cells by ELISA. **(c)** Schematic representation of the experimental design for *in vitro* functional assessment of OSAT. **(d)** Dose-dependent cytotoxic activity of OSAT against B16, MC38, and TC-1 tumor cells at effector-to-target (E:T) ratios of 5:1, 10:1, and 20:1. **(e)** Cytokine secretion profiles (IL-2 and IFN-γ) measured by ELISA at a 20:1 E:T ratio across B16, MC38, and TC-1 tumor models. All graphs represent triplicate experiments. Error bars denote mean ± SEM. Statistical significance was assessed by one-way ANOVA (a and b) or two-way ANOVA (d and e) with tukey’s multiple comparisons test. (*\*P < 0.05, **P < 0.01, ***P < 0.001, ****P <0.0001*; ns = not significant).

### OSAT exhibits superior efficacy and safety in multiple murine tumor models

To investigate the *in vivo* antitumor activity of OSAT compared to traditional TILs, we established syngeneic B16, MC38, and TC-1 tumor models in mice, evaluating both therapeutic efficacy and safety (Fig. 5a). Given the prolonged timeline required for *in vivo* anti-tumor, we confirmed that OSA-GFP delivered by lentivirus maintains high-level sustained expression even after 14 days (Supplementary Fig. 3). In each model, OSAT consistently demonstrated superior antitumor efficacy compared to traditional TILs (Fig. 5b). At 20 days post-treatment, tumors harvested from OSAT recipients were significantly smaller than those from TIL groups (Fig. 5c). Survival analysis further revealed prolonged survival in OSAT-treated mice compared to TIL groups (Fig. 5d).

**Fig. 5.**
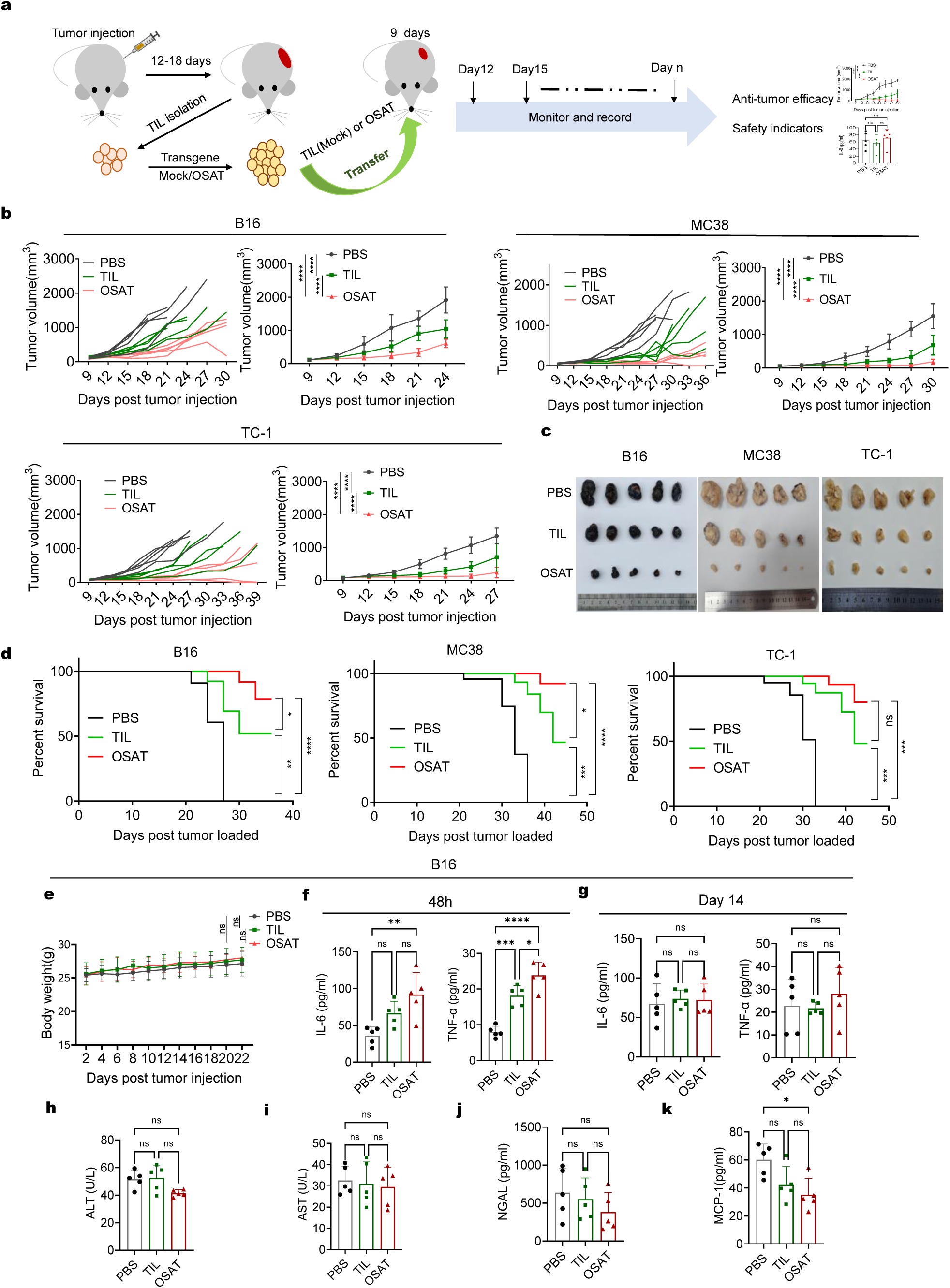
OSAT exhibits remarkable anti-tumor response and advantages in selected tumor models. **(a)** Schematics of experimental design for adoptive transfer treatment in mouse models. **(b)** Tumor growth curves (individuals and groups) of B16/MC38/TC-1 that implanted subcutaneously and subjected to the indicated treatments. **(c)** Photographic images of representative B16/MC38/TC-1 tumors in different groups on day 21 post tumor challenge (n = 5). Representative macroscopic images of excised tumors across treatment groups at endpoint (tumors harvested on Day 21 post-inoculation; n = 5). **(d)** Kaplan-Meier survival curves for B16/MC38/TC-1 tumor-bearing mice treated with PBS, TIL or OSAT. **(e)**Body weight changes in B16 tumor-bearing mice across treatment groups (time-matched with tumor growth monitoring). **(f-g)** ELISA analysis of the cytokine production of IL-6 and TNF-α in the sera of B16 model mice that subjected to the indicated treatment after 48 hours or 14 days. Statistical analysis was performed using non-parametric tests. **(h-k)** Serum biomarkers of hepatic toxicity (ALT, AST) and kidney toxicity (NGAL, MCP-1) at Day 14 post-treatment in the B16 model. The error bars indicate the mean ± the standard error of the mean. Statistical significance was assessed by two-way ANOVA (b and e), by one-way ANOVA (f-k), or by a log rank Mendel–Cox test (d). Each data point indicates an independent donor (n = 5). (*\*P < 0.05, **P < 0.01, ***P < 0.001, ****P <0.0001*; ns = not significant).

Next, we systematically evaluated the safety profile of the treatment regimens. Body weight monitoring showed no significant differences between treatment groups and controls throughout the study (Fig. 5e; Supplementary Fig. 4a, h). Notably, in the TC-1 model, OSAT-treated mice exhibited slight body weight gain compared to controls, likely reflecting reduced tumor burden (Supplementary Fig. 4h). Safety evaluation focused on the risk of CRS *via* serum profiling of IL-6 and TNF-α at 48 hours and 14 days post-treatment, which revealed that cytokine levels in both the TIL and OSAT groups were significantly higher than those in the PBS control group at 48 hours, yet remained within safe and controllable ranges (Fig. 5f-g; Supplementary Fig. 4b-c, i-j). Hepatic and renal function markers (AST, ALT, NGAL, MCP-1) showed no significant differences between groups (Fig. 5h-k; Supplementary Fig. 4d-g, k-n). Collectively, these results demonstrate that OSAT achieve superior antitumor efficacy and confers survival benefits compared to traditional TIL therapy while maintaining acceptable safety profiles.

### Enhanced antitumor efficacy of OSAT combined with PD-1 blockade therapy

We observed that tumors in the B16 tumor model exhibited rapid regrowth following 12 days of OSAT treatment. Thus, we hypothesize that OSAT may progressively upregulate immunosuppressive molecules, thereby attenuating their antitumor activity. To verify this, we analysis the immune checkpoint molecules (PD-1, TIM-3, LAG-3) in TILs isolated from B16 tumors at baseline (day 9) and post-treatment (day 21). Following OSAT treatment, the expression level of PD-1 was significantly elevated compared to the baseline. In contrast, the expression levels of TIM-3 and LAG-3 exhibited no significant changes (Fig. 6a).

**Fig. 6.**
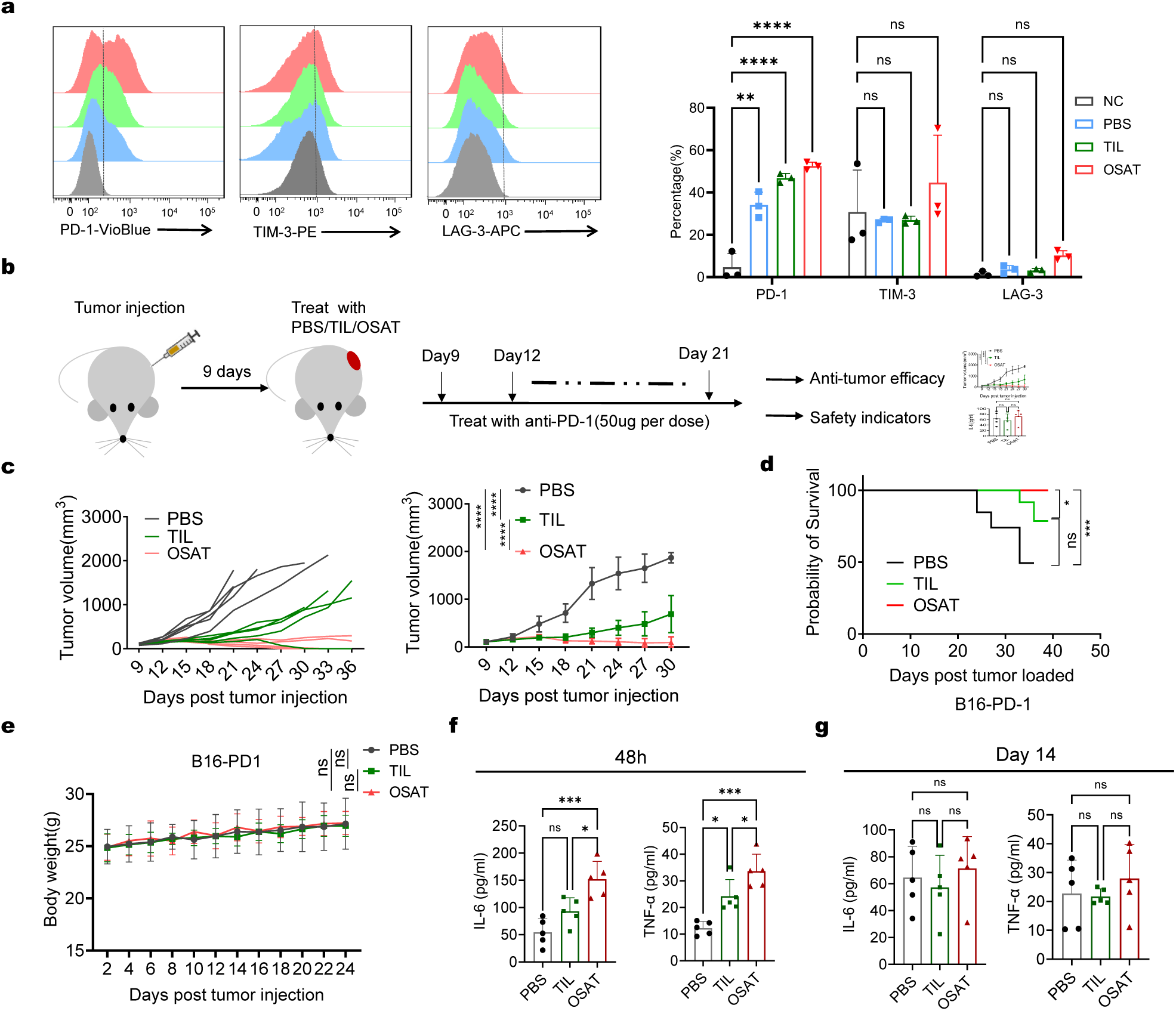
The enhanced antitumor activity of OSAT in combination with PD-1 blockade. **(a)** Comparison of the expression levels of PD-1, TIM-3, and LAG-3 in CD3^+^ T cells before and 12 days after treatment. **(b)** Schematic illustration of the treatment strategy combining TILs with PD-1 blockade. **(c)** Tumor growth curves (individual and group data) of the B16 tumor model mice that received PD-1 blockade and were subjected to the indicated treatments. **(d)** Percent survival curve of B16 tumor-bearing mice that received PD-1 blockade and were subjected to the indicated treatments. **(e)** Body weight changes in the B16 tumor model mice that received PD-1 blockade across different treatment groups (time-matched with the tumor growth monitoring period). **(f-g)** ELISA analysis of the cytokine production of IL-6 and TNF-α in the sera of B16 tumor model mice that received PD-1 blockade and were subjected to the indicated treatments. The analysis was conducted 48 hours or 14 days after treatment. Error bars represent mean ± SEM. Statistical significance was assessed by two-way ANOVA (a, c, and e), by log-rank (Mantel–Cox) test (d) and by one-way ANOVA (f and g). (*\*P < 0.05, **P < 0.01, ***P < 0.001, ****P <0.0001*; ns = not significant).

Next, *in vivo* evaluation of OSAT combined with PD-1 blockade revealed sustained tumor suppression in the B16 model (Fig. 6b-c). Notably, in survival analysis, compared with PD-1 monotherapy, the OSAT plus PD-1 blockade cohort achieved 100% survival by day 42 (Fig. 6d). Furthermore, body weight monitoring confirmed minimal systemic toxicity across groups (Fig. 6e). Serum cytokine profiling at 48 hours and 14 days post-treatment revealed that IL-6 and TNF-α levels were elevated in both the TIL and OSAT groups at 48 hours, yet remained within safe and controllable ranges, indicating acceptable safety profiles (Fig. 6f-g). Moreover, hepatic and renal function markers (ALT, AST, NGAL, MCP-1) showed no significant differences between groups at day 14, confirming acceptable safety (Supplementary Fig. 5a-d). Collectively, OSAT therapy synergizes with PD-1 checkpoint inhibitors to enhance antitumor efficacy while maintaining safety. This combinatorial therapeutic strategy offers a promising approach to overcoming TIL exhaustion and achieving sustained antitumor activity.

## Discussion

An in-depth elucidation of the molecular mechanisms underlying T cell activation serves as the core prerequisite for overcoming the bottlenecks of TIL immunotherapy and enhancing its clinical efficacy^8,30,31^. Given that CARs exhibit high structural relevance to TCRs in terms of function and possess the capacity to precisely regulate signal transduction modules, they are regarded as an ideal simplified model for investigating the signal characteristics of each chain within the TCR-CD3 complex^32,33^. Currently, mainstream CAR designs all incorporate the intracellular domain of the CD3ζ chain, thereby mimicking the signal activation function of the native TCR ^34,35^. Nevertheless, there remains a critical gap in research focusing on the direct genetic modification of the endogenous TCR-CD3 complex: the feasibility of simplifying the T cell activation pathway and efficiently enhancing its anti-tumor activity through this approach has not yet been experimentally validated, which also constitutes a significant barrier restricting the development of TIL therapy toward a more precise and efficient direction. In view of the central role of the CD3ζ chain in TCR signal transduction, and given that previous studies have indicated that CD28 co-stimulatory signals can significantly enhance T cell activation efficiency, this study proposes a strategy of “constructing OSAT by fusing CD3ζ with the CD28-ITAM”. This strategy aims to verify the enhancing effect of single-signal activation on TIL function through the local modification of the endogenous TCR-CD3 complex.

This study provides experimental validation for the hypothesis that engineering TILs to rely on a single activation signal can achieve potent antitumor activity. Our data demonstrate that fusing the CD3ζ and CD28-ITAM bypasses the traditional dual-signaling requirement. Upon mono-signal activation, the OSAT construct exhibits a high degree of similarity in gene expression profiles and functional markers to mock T cells undergoing dual-signal activation. Transcriptomic analysis revealed significant overlap between the OSAT mono-signal activation signature and the dual-signal signature of mock T cells. Surface marker expression (CD25, CD69) reached equivalent levels, indicating that a single integrated signal suffices to drive full activation. *In vitro* cytotoxicity assays and *in vivo* animal experiments demonstrated that OSAT effectively inhibited tumor growth in multiple syngeneic models. Notably, combining OSAT with PD-1 blockade further enhanced therapeutic efficacy, confirming the robustness and clinical translational potential of this approach.

OSAT challenges the classical dual-signal theory of T cell activation while establishing the feasibility of mono-signal activation through functional equivalence and broad antitumor activity across cancer types^36^. By engineering a synthetic signaling domain, we demonstrate that a single integrated signal can substitute for canonical co-stimulation. This aligns with emerging strategies exploring mono-signal activation *via* synthetic receptor design, but uniquely focuses on native TCR diversity in TILs for solid tumor targeting^7,37^. While prior studies have optimized signaling domains in CAR-T cells, our work pioneers the application of mono-signal activation to TILs^38,39^. The inherent tumor-specific heterogeneity and homing capabilities of TILs likely contribute to the observed therapeutic advantages. Evidence suggests that low-affinity T cells drive antitumor responses in humans and mice, and OSA modification may preferentially expand these populations, potentiating long-term efficacy^40^.

However, T cell activation has been linked to T cell exhaustion^41,42^. Our combinatorial approach with PD-1 blockade mitigated this effect, highlighting the need for microenvironment modulation in engineered T cell strategies. The long-term persistence of mono-signal activated T cells warrants further investigation. Despite isolation and CD3⁺ enrichment, TILs retain functional heterogeneity^6,43^. Notably, while OSAT bypasses canonical dual signaling through engineered intracellular domains, a recent study describes a natural two-stage priming process for CD8^+^ T cells, where CXCR3-mediated localization to IL-2 rich niches and CD4-derived paracrine IL-2 drive affinity-based clonal selection^44,45^. Thus, both frameworks emphasize integrating key signaling pathways to optimize activation, proliferation, and functional differentiation for enhanced T cell efficacy.

While this study provides experimental evidence for the feasibility of the OSAT strategy, from the perspectives of research design integrity and clinical translation adaptability, the strategy still has unresolved issues for further optimization and refinement. First, the study only focuses on the fusion modification of the CD3ζ chain with CD28/CD137 and has not systematically investigated other key chains (CD3δ/ε/γ) in the TCR-CD3 complex. It is known that the core structural difference between the CD3ζ chain and CD3δ/ε/γ chains lies in the number of ITAMs in their intracellular domains (the CD3ζ chain contains 3 ITAMs, while each CD3δ/ε/γ chain contains 1 ITAM)^18,46^. Previous studies have confirmed that a single ITAM is sufficient to support CAR-T cells in exerting anti-tumor functions and may even reduce the risk of cytokine release syndrome^32,33,47^. Future research needs to systematically explore the mono-signal activation potential of CD3δ/ε/γ chains, and compare the activation efficiency, cytokine secretion characteristics, and side effect risks after modification of different CD3 chains. The value of this research direction lies in identifying optimal modification targets in the TCR-CD3 complex, thereby providing a basis for developing “highly effective and low-toxic” OSAT variants. Second, the validation system of this study only includes primary human T cells and immunocompetent mouse models and does not involve humanized tumor microenvironment models. Meanwhile, it has not analyzed the differences in OSAT modification effects on TILs from different individuals, yet the inherent heterogeneity of individual TILs is a key factor affecting the efficacy of TIL therapy^48,49^. Future studies need to validate OSAT’s efficacy in humanized models, expand samples to analyze its effects on TILs from different individuals, and identify sensitive T cell subsets—this direction can support OSAT’s clinical translation and personalized TIL therapy. Third, although this study confirms that OSAT combined with PD-1 blockade can alleviate T cell exhaustion, it remains unclear whether OSAT itself affects the T cell exhaustion process through specific molecular pathways. Furthermore, the long-term *in vivo* persistence of mono-signal activated T cells also lacks systematic evaluation, such as memory cell differentiation potential and sustained effector cytokine secretion capacity^50,51^. Future research should use techniques such as proteomics and single-cell sequencing to elucidate the molecular mechanisms by which OSAT regulates T cell exhaustion and assess the *in vivo* persistence and memory function of OSAT-modified T cells. This can provide a mechanistic basis for further optimizing the OSAT strategy and improving long-term therapeutic efficacy.

In summary, this work introduces a novel engineering strategy for TILs, generating mono-signal activated TILs with enhanced antitumor activity in preclinical models. This approach addresses critical limitations in current TIL therapy and may facilitate broader application in solid tumor treatment.

## Methods

### Generation of OSA constructs

DNA fragments encoding human/mouse CD3ζ, CD28, and CD137 were chemically synthesized (Sangon Biotech). Fragments were cloned into pLVX-Puro for lentiviral packaging using Gibson assembly^52^. Linearized/dephosphorylated vectors and PCR-amplified inserts were assembled following standard protocols. OSA fragments were also TA-cloned into pGEM-T Easy (Promega) for gene-editing donor use and ligated into modified pMC plasmids containing the APOB 3’ MAR element^22^. The *CMV* promoter was replaced with the human *EF1α* promoter. Final constructs were validated by Sanger sequencing (Sangon Biotech).

### Cell lines

All cell lines were obtained from the American Type Culture Collection (ATCC) and authenticated by short tandem repeat (STR) profiling within the last 6 months prior to experimentation. The human T cell cancer cell line Jurkat E6-1 and mouse T cell cancer cell lines EL-4, as well as the mouse colon cancer cell line MC38 and lung cancer cell line TC-1, were cultured in RPMI-1640 medium (Gibco, 11875500) supplemented with 10% heat-inactivated FBS (Gibco, 10099141), 100 U/ml penicillin-streptomycin (Sangon Biotech, A506002). HEK293T cells (ATCC, CRL-1573) were cultured in DMEM (Gibco, 11965092) supplemented with 10% heat-inactivated FBS, 100 U/ml penicillin-streptomycin. B16 cells (ATCC, CRL-6322) were cultured in the same DMEM medium with the same supplements. Both cell lines were incubated at 37 °C in a 5% CO₂ humidified incubator. Cells were passaged when they reached 70-80% confluence using 0.05% trypsin-EDTA (Gibco, 25300054) for detachment.

### Lentivirus packaging and transduction

HEK293T packaging cells were seeded at a density of 3 × 10^6^ cells per 10-cm dish in DMEM supplemented with 10% FBS and incubated at 37℃ in a humidified atmosphere with 5% CO2. Transfection was carried out using Lipofectamine 2000 Reagent (Thermo Fisher Scientific, 11668019) following the manufacturer’s protocol: a mixture of 3 μg of OSA-expressing plasmids and 9 μg of lentiviral packaging plasmids, pMD2.G (Addgene, 12259) for envelope and psPAX2 (Addgene, 12260) for packaging components, with total DNA adjusted to 12 μg, was incubated at room temperature for 20 min and added drop-by-drop to the cells. 48 hours post-transfection, supernatants with lentiviral vectors were harvested, filtered through a 0.45 μm filter (Millipore, SLHV004SL), and concentrated with a 10% sucrose solution by ultracentrifugation at 100,000g for 4 h at 4℃ in a Beckman Coulter SW32Ti rotor. Virus preparations were titrated using HEK293T cells as targets: serial dilutions were added to HEK293T cell cultures, and 72 h later, the number of transduced cells was determined by detecting GFP expression *via* flow cytometry. For Jurkat cell transduction, Jurkat cells were seeded at a density of 2 × 10^6^ cells per well in 6-well plates in RPMI-1640 medium with 10% FBS and 1% penicillin-streptomycin, and concentrated lentiviral vectors were added at a multiplicity of infection (MOI) of 48 h later, 1 μg/ml puromycin (Sigma Aldrich, P8833) was added for two-week selection. All procedures were performed in a biosafety level 2 (BSL-2) laboratory following standard safety protocols for handling lentiviral vectors.

### ELISA analysis

Cytokine secretion profiles of interleukin-2 (IL-2), interferon-γ (IFN-γ), and tumor necrosis factor-α (TNF-α) from human primary T cells, Jurkat cells, murine T cells, and EL-4 cells were quantified using sandwich ELISA kits. The kits used were from Thermo Fisher Scientific: for human samples, IL-2 (BMS221-2), IFN-γ (KHC4021), TNF-α (KHC3014C); for mouse samples, IL-2 (BMS601), IFN-γ (BMS6027). For the analysis of serum IL-6 (Beyotime Biotechnology, PI326), TNF-α (Beyotime Biotechnology, PT512), hepatic markers alanine aminotransferase (ALT, Abcam, ab105134) and aspartate aminotransferase (AST, CUSABIO, CSB-E12649m), and renal biomarkers neutrophil gelatinase-associated lipocalin (NGAL, Bio-Swamp, ZN2689) and monocyte chemoattractant protein-1 (MCP-1, Lianke Biotech, EK287HS-96), assays were carried out following the manufacturer-recommended protocols. Absorbance measurements were conducted on a BioTek Synergy H1 microplate reader (Agilent) with dual-wavelength correction at 450 nm/570 nm. Data were normalized to standard curves generated using a 4-parameter logistic fit, with an R² value > 0.99.

### Identification of intracellular calcium flux

For intracellular calcium flux identification, cells were loaded with Fluo-8 AM (Abcam, ab142773) (100 μl, 4 μM) in Hanks’ Balanced Salt Solution (HHBS) supplemented with 20 mM Hepes, and incubated at 37 °C with 5% CO₂ for 1 h in the dark, followed by two washes with HHBS. Cells were subsequently stimulated with 1 ug/ml anti-CD3 antibody (Sino Biological, OKT3, GMP-10977) /anti-CD28 antibody (Sino Biological, CD28.2, GMP-11524) alongside various treatments, then analyzed using a fluorescence microplate reader. Timepoint values represent means from three independent central views across triplicate wells. Fluorescence intensity analysis utilized maximum group values (MAX) as baseline, with real-time MFI (Mean Fluorescence Intensity) expressed as percentage change relative to baseline.

### Western blots

Jurkat or OSA-Jurkat cells were stimulated with 1 µg/ml of anti-CD3 and/or anti-CD28 antibodies. After 1 hour, the cells were lysed using RIPA lysis buffer (Thermo Fisher Scientific, 89900) containing sodium orthovanadate and protease inhibitor (Thermo Fisher Scientific, 78430), incubating on ice for 15 minutes. The lysates were then subjected to high-speed centrifugation to remove debris. Samples were combined with 5 × SDS loading buffer and heated at 95°C for 10 minutes. The protein samples were loaded onto gels and electrophoresed at 120 V for 90 minutes. Following this, proteins were transferred to nitrocellulose membranes using transfer buffer at 100 V for 35 minutes. The membranes were blocked with nonfat dry milk for 1 hour and subsequently washed with Tris-buffered saline containing 0.1% Tween-20 (TBST). After three washes with TBST, the membranes were incubated overnight with a primary antibody, followed by a 1-hour incubation with a horseradish peroxidase (HRP)-conjugated secondary antibody. The membranes were washed three times with TBST and then probed using enhanced chemiluminescence solution (Coolaber, SL1351). The antibodies utilized in this experiment included GAPDH (Proteintech, 10494-1-AP, 1:20000), NF-κB (p65) (Abcam, ab28856, 1:3000), and AP-1 (JUN-C) (Abbkine, ABP50261, 1:3000).

### RNA-seq analysis of OSA-Jurkat cells

Total RNA from OSA-Jurkat cells was isolated using Trizol Reagent. After assessing its quality with a NanoDrop spectrophotometer, 3 μg of RNA was used to synthesize cDNA with the SMARTer Ultra Low Input RNA Kit. Sequencing libraries were prepared using the Nextera XT kit (Illumina). The process included mRNA purification, fragmentation, cDNA synthesis in two steps, end-modification, adapter ligation, size-selection (400-500 bp) with the AMPure XP system, and PCR enrichment (15 cycles). The libraries were quantified using the Agilent high-sensitivity DNA assay on a Bioanalyzer 2100 system. Sequencing was carried out on a P3 flowcell-NextSeq2000 platform. Raw sequencing data was processed with fastp (v0.22.0) to obtain clean reads, which were then mapped to the reference genome using HISAT2 (v2.1.0). Gene expression was analyzed by HTSeq (v0.9.1) and normalized with FPKM/TPM^34^. Differential gene expression was determined by DESeq2 (v1.38.3) with criteria of ∣log2FoldChange∣ > 1 and *P < 0.05*. All analyses were performed on the Personalbio Genes Cloud platform.

### CRISPR–Cas9 knockin

CRISPR-Cas9-mediated knockin was performed using a modified approach based on previously published methods^27^. Briefly, sgRNA targeting the CD3Z locus (guide sequence: *5′-TTTCACCGCGGCCATCCTGC-3′*) was designed and cloned into the BbsI restriction site of the PX330 vector (Addgene, 42230) *via* Gibson assembly. The donor sequences, synthesized by Genewiz, were subsequently cloned into the P-EASY plasmid. A total of 10 µg of Cas9 protein and 10 µg of donor DNA were used for electroporation of 2 million T cells using the preset DS104 program of the 4D-Nucleofector™ system (Lonza). Electroporation was conducted according to the manufacturer’s instructions. After electroporation, cells were resuspended in pre-warmed RPMI-1640 medium containing 10% FBS.

### Primary human T cell isolation, activation and expansion

Whole blood obtained from healthy donors (BER-YXY-2024023) PBMCs were isolated from whole blood using density gradient centrifugation with Ficoll-Paque PREMIUM (1.077 g/mL)^53^. Briefly, blood was diluted 1:2 with saline, layered over separation medium, and centrifuged at 750 × g for 15 min (brake disabled). The PBMC layer was collected, washed twice in PBS, and centrifuged at 200 × g for 5 min. Autologous plasma was heat-inactivated (56°C, 30 min) and stored at 4°C for subsequent use. CD3^+^ T cells were isolated *via* magnetic-activated cell sorting (MACS) using CD3 MicroBeads (MiltenyiBiotec). Cells were incubated with CD3 MicroBeads (20 μL per 1 × 10⁷ cells) in sorting buffer (DPBS + 0.5% BSA + 2 mM EDTA) at 4°C for 15 min, washed, and passed through a pre-rinsed LS column. The CD3^+^ fraction was eluted after removing unbound cells. For activation and expansion, CD3 antibodies (1 μg/mL) were immobilized on 6-well plates overnight. Purified T cells were cultured in RPMI-1640 medium supplemented with 10% FBS, CD28 antibody (1 μg/mL), and IL-2 (200 IU/mL) at 2.5 × 10⁵ cells/mL. Cultures were maintained in a 5% CO₂ incubator for 2–5 days, achieving about 5-fold expansion. The purity and activation status of T cells were evaluated by flow cytometry.

### Flow cytometry analysis

Flow cytometry was utilized to assess the efficiency of gene editing or gene introduction mediated by the OSAT molecule. CD3⁺ T cells harvested at distinct time points were stained with fluorescently-conjugated antibodies specific for human CD25 (PE, Miltenyi, 130-095-942), human CD69 (APC, Milteny, 130-095-891), mouse anti-CD3 (FITC, Miltenyi, 130-092-277), mouse anti-PD-1 (VioBlue, Miltenyi, 130-121-729), mouse anti-TIM-3 (PE, Miltenyi, 130-116-816), and mouse anti-LAG-3 (APC, Miltenyi, 130-120-435), followed by flow cytometric analysis. All data were processed using FlowJo software.

### T-cell cytotoxicity assay

T cells and target cells were counted and mixed in a 1:1 ratio, with 2 × 10^5^ cells added to each well of a 24-well plate. After a 24-hour co-culture period, the cells were harvested and washed three times with flow cytometry staining buffer. The samples were subsequently stained with Annexin V-APC and DAPI (Elabscience, E-CK-A258) for 15 minutes at room temperature. FlowJo software was utilized for data analysis. The T cell-mediated killing rate (%) was determined by calculating the percentage of Annexin V-positive cells among the tumor cells (FITC^+^) co-cultured with T cells, subtracting the percentage of Annexin V-positive tumor cells cultured alone. The formula for the T cell-killing rate is as follows: T cell-killing rate (%) = Annexin V positive rate (%) of tumor cells co-cultured with T cells-Annexin V positive rate (%) of tumor cells cultured alone. The assessment of Annexin V positivity was conducted in accordance with established methodologies^54^.

### Murine tumor mouse models

Female C57BL/6 mice, 6-8 weeks old, were obtained from Charles River Laboratory (Beijing, China) and randomly allocated to either the experimental or control group. Mice were housed in specific pathogen-free animal facilities at the Hebei University of Engineering, maintained on a 12h light–dark cycle. The housing temperature was controlled within a range of 21-24 °C, and relative humidity was maintained between 25% and 61%. Mice were provided with diet. All animal experiments were carried out in strict accordance with the guidelines and regulations of the Hebei University of Engineering Animal Care and Use Committee, Hebei, China.

The mice were first shaved, and then underwent subcutaneous injection. For the injection of tumor cells, three types of cells were used: B16 cells at a density of 2 × 10⁵ cells per 0.1 ml per mouse, MC38 cells at 5 × 10⁵ cells per 0.1 ml per mouse, and TC-1 cells at 5 × 10⁵ cells per 0.1 ml per mouse. During the experimental process, the tumor volumes were regularly evaluated. Specifically, the size of the tumor was measured in two dimensions using a digital caliper every 2-3 days. The tumor volume was calculated by the formula: Volume = (length × width²)/2. In some experiments, such as those related to anti-tumor efficacy, if the tumor reached 15 mm in its longest axis, or developed ulceration or necrosis, it was considered as an endpoint, similar to the definition in previous studies.

### Mouse CD3^+^ T cell isolation

Tumors were excised 12-18 days post tumor engraftment. Fresh tumor tissues were carefully washed with appropriate buffer solutions and then cut into approximately 1 mm³ pieces. To facilitate tissue dissociation, the samples were incubated with a mixture of DNase, collagenase (such as collagenase IV at a concentration of 1 mg/ml), and hyaluronidase. This digestion process was carried out at 37 °C in a shaking incubator set at 200 rpm for 30 minutes. After digestion, the cell suspension was filtered through a 70 μM cell strainer to remove large debris. Subsequently, density gradient centrifugation, for example, using Percoll, was performed to separate lymphocytes. The white cell layer, which is enriched with TILs and located between 75% and 100% separation liquid layers, was carefully collected. For the isolation of CD3^+^ T cells, positive selection was carried out using mouse CD4/CD8 (TIL) MicroBeads (MiltenyiBiotec, 130-095-249), following the detailed manufacturer’s instructions precisely. When isolating CD3^+^ T cells from PBMCs, density gradient centrifugation was first used to separate lymphocytes. Then, CD3^+^ T cells were isolated from the white cells using the same mouse CD4/CD8 (TIL) MicroBeads according to the manufacturer’s protocol. The purified CD3^+^ T cells were amplified in RPMI-1640 medium supplemented with 10% FBS and a high dose of mouse IL-2 (eBioscience, 14-8021-82).

### CCK8 assay

The effector cells include TILs and OSAT, and the corresponding tumor cell lines were used as target cells. A 96-well plate was prepared with control wells containing RPMI-1640 complete medium supplemented with 10% FBS, as well as experimental and control wells. Subsequently, 100 μl of target cells (2 × 10⁴/ml) were seeded 24 hours in advance, and 100 μl of effector cells (1 × 10⁵/ml, 2 × 10⁵/ml, 4 × 10⁵/ml) were added to the experimental wells. The plate was then incubated at 37°C with 5% CO2 for 24 hours. Afterward, all supernatants were aspirated, and the wells were washed twice with PBS before adding 100 μl of complete medium. 12 hours later, 10 μl of CCK-8 solution (MCE, HY-K0301) was added to each well. Following a 4-hour incubation, the *in vitro* anti-tumor activity of TILs was assessed using the CCK-8 method according to the manufacturer’s instructions.

### *In vivo* murine tumor experiments

To assess the *in vivo* anti-tumor activity of OSAT compared to traditional TIL, we conducted experiments using three tumor models over two time periods. In the first period, tumor tissue was removed, and CD3^+^ TIL cells were extracted 12-18 days after tumor formation. After amplification with IL-2, half of the TIL cells were introduced to the OSA plasmid through electroporation, referred to as OSAT, while the remaining half remained as traditional TIL. These cells were then injected back into the tumor model mice *via* tail vein injection (1 × 10^7^ cells/mice). Anti-PD-1 antibody (RMP1-14, BioXcell) was injected intraperitoneally in the corresponding groups at day 12, 15, 18 and 21 after tumor cell inoculation. Tumor volume was measured every three days using a Vernier caliper, and survival time was recorded. Tumor volume was calculated using the formula: volume = (length × width^2^)/2.

### Statistical analysis

Statistical analysis was performed using GraphPad Prism software (v10.3.1). Data are presented as mean ± standard deviation unless otherwise stated. Statistical tests included one-way or two-way ANOVA with tukey’s multiple comparisons test and log-rank (Mantel-Cox) test. Differences were considered statistically significant when *P* < 0.05. All statistical tests were two-sided, and sample sizes were determined based on prior experience and power analysis to ensure sufficient statistical power to detect meaningful differences.

## Data and materials availability

All other data are available in the maintext or the Supplementary Information. Source data are provided with this paper. Transcriptome data have been deposited in the NCBI Sequence Read Archive (BioProject ID: PRJNA1237508).

## Acknowledgments

We thank the facility support of Gene Editing Research Center, Hebei University of Science and Technology.

## Author contributions

Yongqiang Wu and Xueshuai Ye conceived the project, designed all experiments. Yongqiang Wu, Xueshuai Ye, Yufeng Zhu, Shengchao Liu, Jiabao Song and Yutong Zhang performed the experiments. YongqiangWu, Xueshuai Ye, Qichen Yuan, Dejing Kong and Lianmei Zhao analyzed the data. Yongqiang Wu, Fei Liu and Lianmei Zhao wrote the manuscript with help from all authors.

## Ethics statement

This study was approved by the Ethics Committee of The Affiliated Hospital and School of Clinical Medicine, Hebei University of Engineering (Approval Number: BER-YXY-2024023).

## Supplementary Figures

**Fig. S1:**
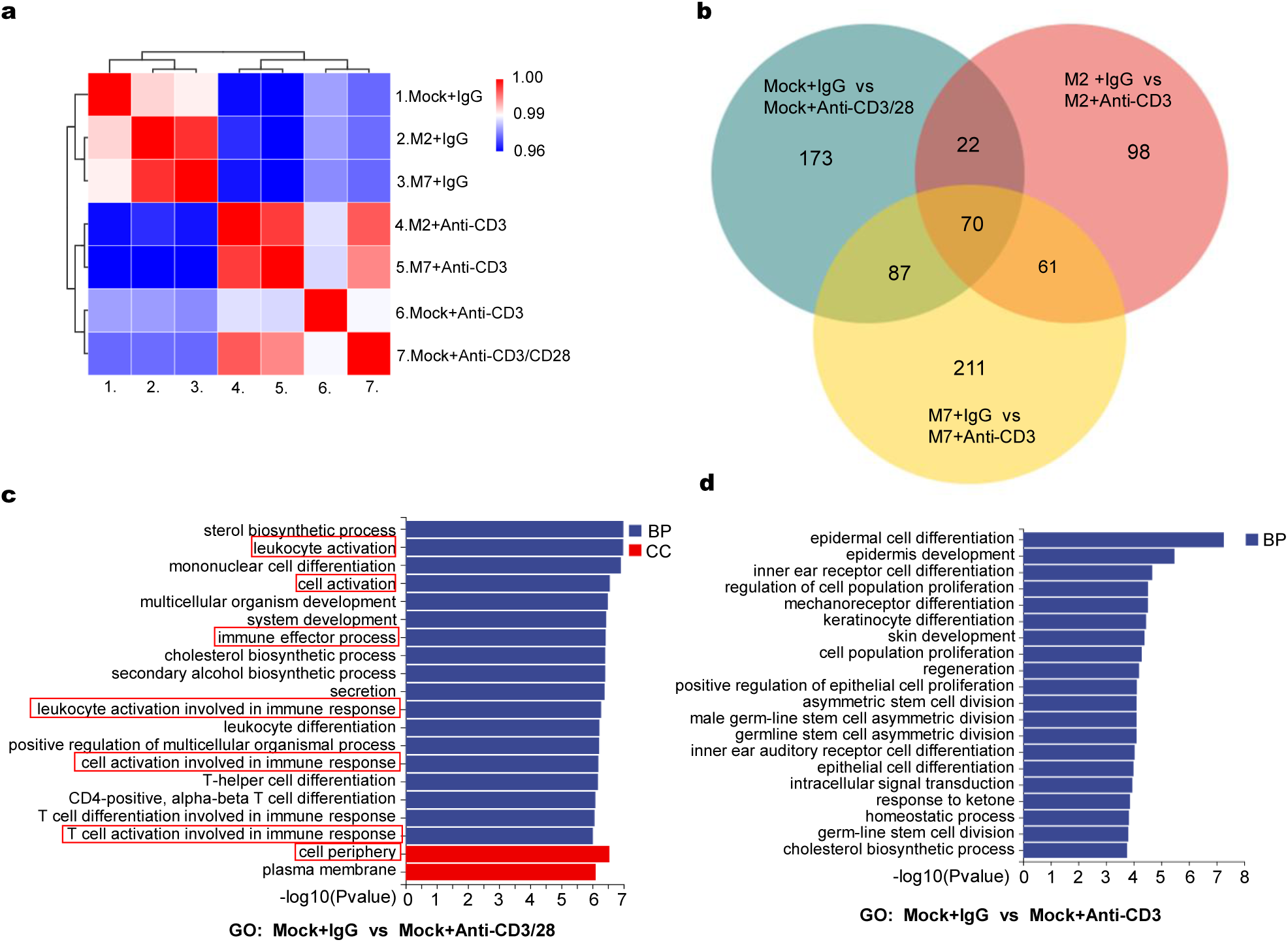
Transcriptomic profiling validated functional equivalence. **(a)** Principal component analysis of global gene expression profiles in WT-Jurkat vs. OSA-Jurkat cells. Cells were stimulated for 12 hours, and transcriptomic data were normalized using Z-score transformation. **(b)** Venn diagram of differentially expressed genes. **(c-d)** Gene Ontology (GO) enrichment analysis comparing Mock + IgG vs. Mock + Anti-CD3/28 and Mock + IgG vs. Mock + Anti-CD3 Jurkat cells. The top 20 most significantly enriched GO terms for differentially expressed genes are shown. GO enrichment analysis was performed using the cluster Profiler R package. Differentially expressed genes were annotated to GO terms, and enrichment was calculated *via* hyper geometric distribution. Statistical significance was defined by a P-value threshold of < 0.05, identifying GO terms overrepresented in differential gene sets relative to the genomic background. This analysis characterized the primary biological functions associated with differentially expressed genes.

**Fig. S2:**
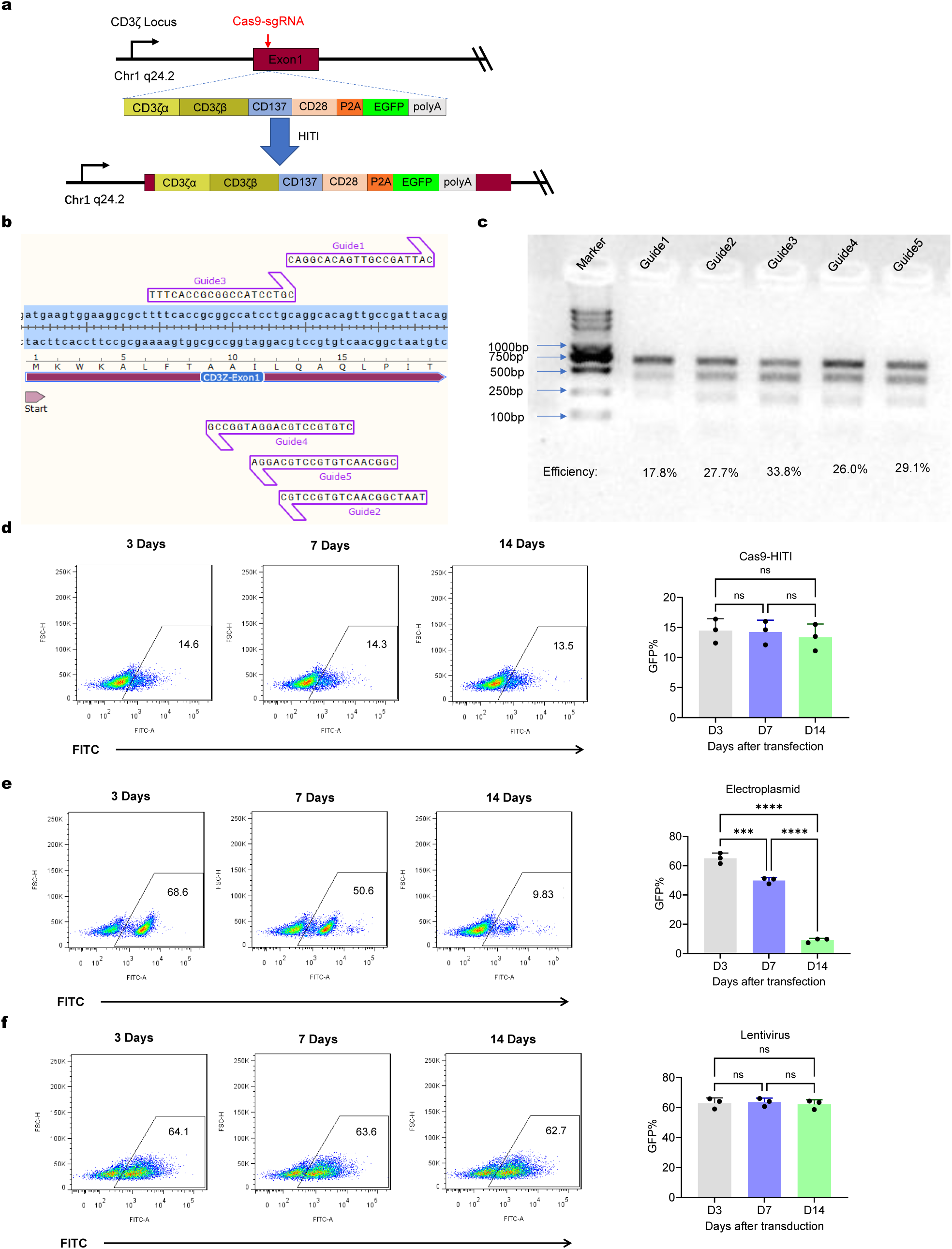
Design and characterization of Cas9-HIHI genome editing and expression persistence comparison of Cas9-HITI, electroplasmid, and lentivirus systems. **(a)** CRISPR-Cas9/HITI-mediated integration of the OSA gene and reporter gene (eGFP) into the CD3ζlocus. **(b)** Schematic representation of the genomic locations of the five single-guide RNAs (sgRNAs) targeting the *CD3Z* gene. **(c)** Detection of indels at the *CD3Z* exon 1 locus using Cas9-sgRNA complexes with five different guides. PCR amplification was performed using primers CD3Z-F (GCGAATTTCTTGGCCCTGTC) and CD3Z-R (GTTTCTCTACCCTGGGCCTG). Annealed PCR products were digested with T7 Endonuclease I (New England Biolabs) for 30 min, resolved by 1.5% agarose gel electrophoresis, and quantified by band intensity analysis. Indel efficiency was calculated using ImageJ software *via* the formula: 100 × [1 − √(1 − (b + c)/(a + b + c))], where *a* denotes the integrated intensity of the undigested PCR fragment and *b*/*c* represent the intensities of cleavage products. **(d-f)** Persistence of plasmid expression following Cas9-HITI,electroplasmid and lentivirus. Error bars represent mean ± SEM. Statistical significance was determined by one-way ANOVA with tukey’s multiple comparisons test (*\*P < 0.05, **P < 0.01, ***P < 0.001, ****P <0.0001*; ns = not significant).

**Fig. S3:**
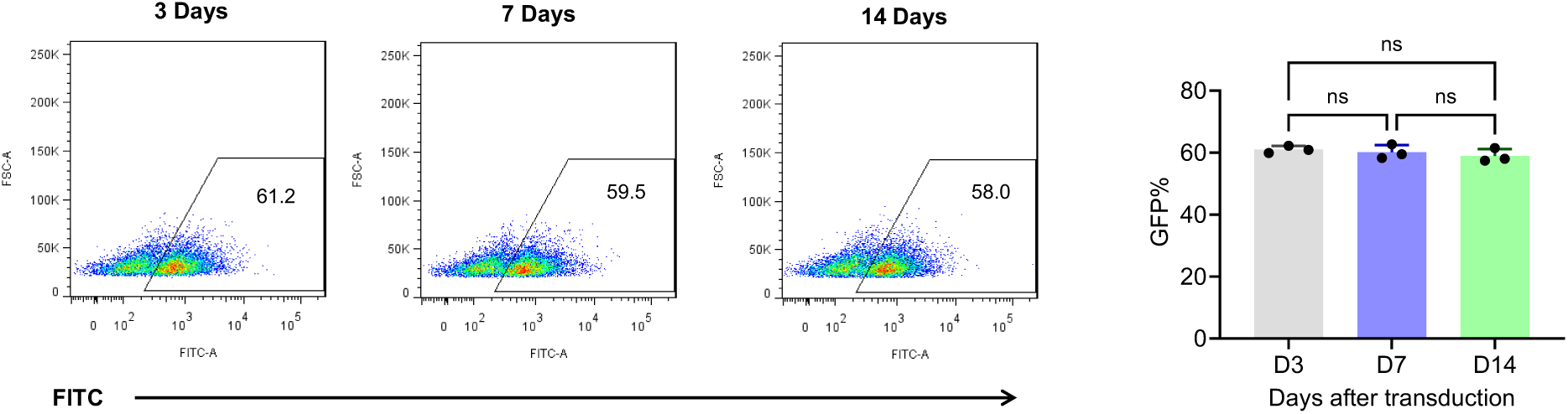
Duration of OSAT molecule expression in murine cells delivered by lentivirus. Efficiency and persistent expression of OSAT molecules delivered into the EL-4 murine T cell line *via* lentiviral infection. Error bars represent mean ± SEM. Statistical significance was determined by one-way ANOVA with tukey’s multiple comparisons test *(*P < 0.05, **P < 0.01, ***P < 0.001, ****P <0.0001*; ns = not significant).

**Fig. S4:**
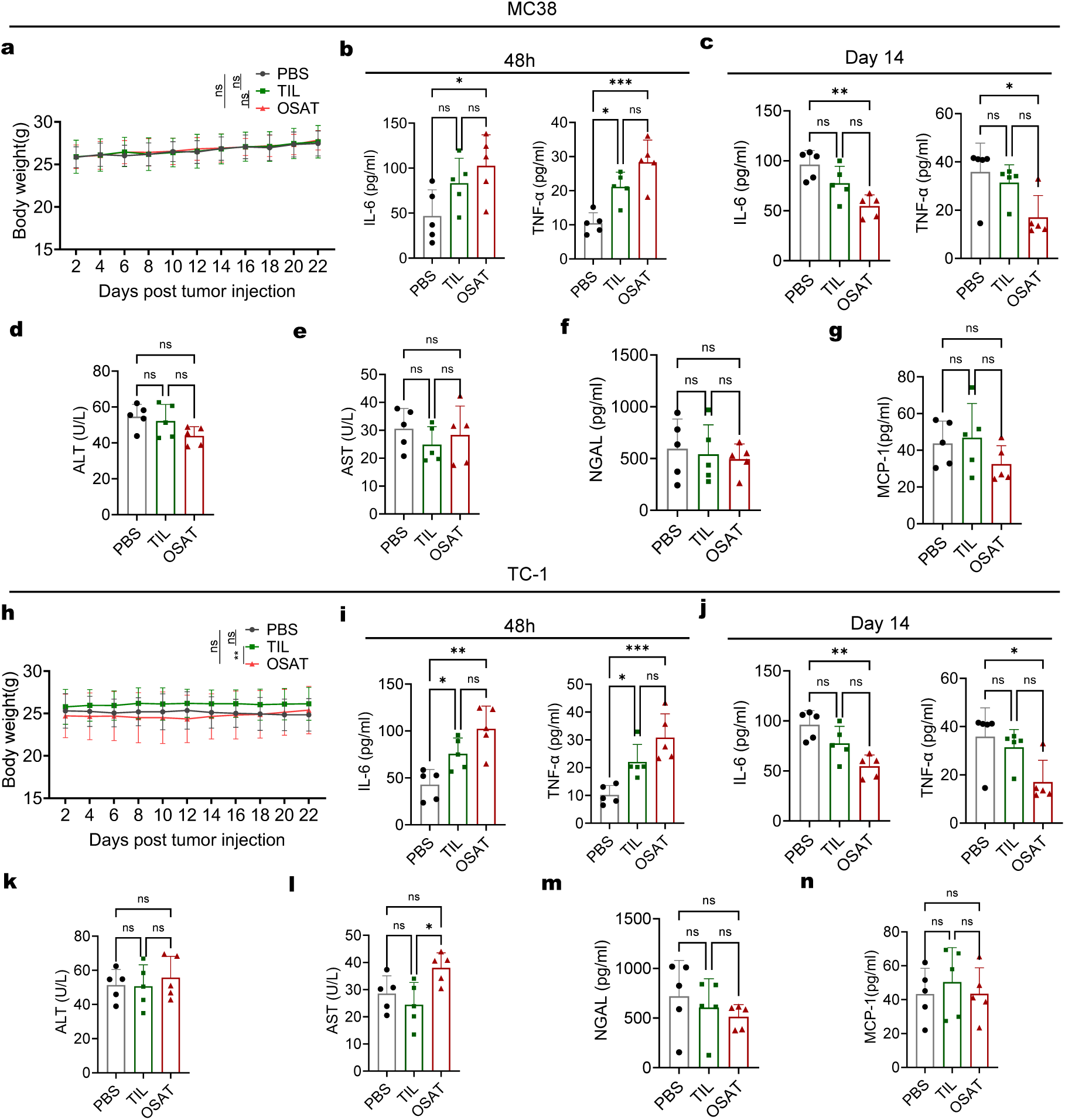
OSAT demonstrates remarkable anti-tumor response and advantages in MC38 and TC-1 models *in vivo*. **(a)** Body weight changes in MC38 tumor-bearing mice across treatment groups (time-matched with tumor growth monitoring). **(b-c)** ELISA analysis of the cytokine production of IL-6 and TNF-α in the sera of MC38 model mice that subjected to the indicated treatment after 48hours or 14days.Statistical analysis was performed using non-parametric tests. **(d-e)** Serum biomarkers of hepatic/kidney toxicity (ALT, AST) and inflammation (NGAL, MCP-1) at Day 14 post-treatment in the MC38 model. **(h)** Body weight changes in TC-1 tumor-bearing mice across treatment groups (time-matched with tumor growth monitoring). **(i-j)** ELISA analysis of the cytokine production of IL-6 and TNF-α in the sera of TC-1 model mice that subjected to the indicated treatment after 48 hours or 14 days. Statistical analysis was performed using non-parametric tests. **(k-n)** Serum biomarkers of hepatic toxicity (ALT, AST) and kidney toxicity (NGAL, MCP-1) at Day 14 post-treatment in the TC-1 model. The error bars indicate the mean ± the standard error of the mean. Statistical significance was determined by one-way or two-way ANOVA with tukey’s multiple comparisons test (\**P < 0.05, **P < 0.01, ***P < 0.001, ****P <0.0001*; ns = not significant).

**Fig. S5:**
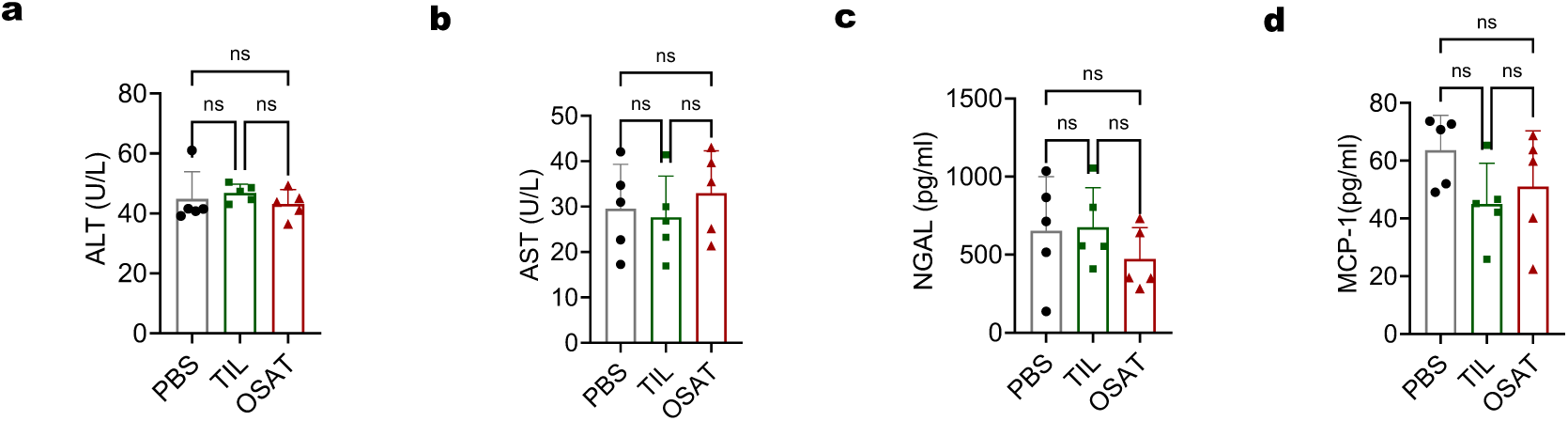
Hepatic and renal safety biomarkers following combination with PD-1 blockade. **(a-d)** Serum biomarkers of hepatic toxicity (ALT, AST) and renal toxicity (NGAL, MCP-1) in the TC-1 model at day 14 post-treatment. Error bars represent mean ± SEM. Statistical significance was determined by one-way ANOVA with tukey’s multiple comparisons test (*\*P < 0.05, **P < 0.01, ***P < 0.001, ****P <0.0001*; ns = not significant).

